# Incomplete adaptation to surface movement during hand reaching

**DOI:** 10.1101/2024.12.18.629126

**Authors:** Priscilla Balestrucci, Matteo Bianchi, Colleen P. Ryan, Giulia Daniele, Alice Flamini, Fanny Valente, Francesco Lacquaniti, Alessandro Moscatelli

**Affiliations:** Laboratory of Neuromotor Physiology, Santa Lucia Foundation IRCCS, Rome, 179; Department of Systems Medicine and Centre of Space Biomedicine, University of Rome Tor Vergata, Rome, 00133; Department of Systems Medicine, University of Rome Tor Vergata, Rome, 00133; Research Centre E. Piaggio and Department of Information Engineering, University of Pisa, Pisa, 56122

## Abstract

Humans can effectively adapt to changes in the environment to maintain adequate motor performance in a vast range of situations. However, residual errors tend to persist when strong *a priori* assumptions about the statistical regularities of the environment are violated. In our study, we challenged the expectation that inanimate objects are usually at rest. To this end, we used a robotic interface to move a plate over which participants slid their finger while reaching towards a target. We found limited evidence of adaptation after prolonged exposure to this perturbation, and only when visual feedback about hand position was provided. Although participants were aware of the motion of the contact surface, explicit knowledge about its direction was limited. Our results provide important insights on the limits of adaptation to motion perturbation in the somatosensory system, which can inform the design of technology applications such as haptic interfaces and collaborative robots.

## 1. Introduction

The brain has the remarkable ability to adapt motor behavior when facing a vast range of changes in the surrounding environment and in one’s own body ^1,2,3^. Reaching and throwing movements are well-studied instances of this adaptability, where rapid adaptation occurs in response to a variety of changing environments ^4,5,6^. Different types of perturbation are employed to study adaptation in motor control. For instance, researchers may change the mapping between performed reaching movements and visual information. In many studies, this was achieved by introducing a linear shift ^7,8^ or angular rotation ^9^ between the actual arm position and its visual representation. In other tasks, experimenters evaluate adaptation following modifications to limb dynamics. This involves, for example, introducing viscous ^10,11,12^ or inertial forces ^13,14^ perturbing the mechanical features of the reaching movement. In general, the introduction of a perturbation leads to systematic motor errors, in the form of increased distance from the target at the end of the reaching movement or deviation of the reaching trajectory from the intended path. With consistent exposure to the perturbation, typical patterns of adaptation generally arise, where the decrease of error over time follows an exponential decay towards baseline performance; moreover, when the perturbation is removed, there is an error increase in the opposite direction than the one occurring in the presence of the perturbation (aftereffect), followed again by an exponential decay and fast return to baseline performance.

Adaptation does not usually occur or is limited when the perturbation violates a strong prior about our body or the world around us. The central nervous system relies on the statistical regularities of the environment in perceptual and motor tasks ^15,16,17^. Thus, humans assume *a priori* the presence of phenomena such as the pull of Earth gravity ^18,19,20^, a source of light from above ^21^, or frictional forces ^22,23^. These assumptions contribute to interpret incoming sensory stimuli and to predict future states of external objects and of our own body by means of feed-forward internal models. If, on the one hand, this approach allows for rapid and effective motor control within the bounds of familiar and recurrent conditions, on the other hand it can give rise to perceptual illusions ^24,25^ and systematic motor errors ^18,26^ once a strong prior is violated. Such is the case, for instance, of catching and throwing movements under modified gravity conditions ^18,27^. In their study, Zago and Lacquaniti (2005) propose that the lack in motor adaptation in this altered environment is due to our limited ability to update the internal model of the strong prior–in this case, gravity–through practice.

While the expectation of gravity is seldom violated in everyday life, other environmental conditions can more easily challenge the *a priori* assumptions made by the brain. For example, human observers assume that self-motion of inanimate objects is unlikely ^28,25,29^. Given this assumption, referred to as static or slow-motion prior, motion relative to our sensory sheets, the skin for touch or the retina in vision, is more likely to be interpreted as our body or our gaze moving with respect to a static object rather than inanimate objects moving relative to us. This prior holds true in a broad spectrum of daily situations, but it can be challenged in real-world scenarios—such as the common experience of feeling self-motion while standing on a train platform next to a moving train—as well as in laboratory settings ^28^.

The static prior plays a crucial role in interpreting somatosensory stimuli when interacting with physical objects, e.g. during reaching and grasping, or when manipulating them to comprehend their properties. In fact, when actively sliding a finger on a moving surface, the estimation of motion of the surface is biased, with a tendency to perceive it as world-stationary ^22^. In the motor domain, systematic errors in reaching movements arise when the finger slides over a surface in motion ^30^. In other words, slip motion perceived at the skin is interpreted as a consequence of hand movement, even when it is due to surface movement.

With the rising availability of technology such as immersive virtual and mixed reality environments and collaborative robots, the prior of stationarity applied to inanimate objects can be easily challenged in simple contexts, and even exploited for innovative applications, for example in the control algorithms for haptic retargeting ^31^. Thus, exploring the limitations imposed by interactions with these types of technology is both intriguing and necessary. In particular, it becomes important to investigate whether humans can effectively cope with and adapt to systematic violations of the static prior.

To this end, we investigated whether human participants are able to correct their movements across trial repetitions to account for the displacement of a contact plate in a reaching task. More specifically, we evaluated whether prolonged exposure to a novel condition of surface motion induces adaptive changes in motor control. We asked participants to slowly slide a finger over a plate to reach a virtual target, while their hand remained hidden from view (Figure 1 A,B). After a series of 20 trials in which the contact plate remained in a fixed position (baseline block), a perturbation was introduced that caused the plate to move with respect to the finger (perturbation block). The position of the plate was continuously updated via the robotic interface illustrated in Figure 1 A-B proportionally to the displacement of the fingertip sliding above it, up to a gain factor (as detailed in the Methods section and in ^32^; ^33^). This caused a movement of the plate orthogonal to hand motion, and with a speed relative to the finger dependent on the value of the gain factor. Such perturbation was maintained consistently for 125 trials, and removed in 10 catch trials interspersed within the last 90 trials of the perturbation block. The contact plate was stationary in the last 20 trials of the experimental procedure (washout block). The perturbation applied to the plate caused the slip motion perceived at the fingertip to be inconsistent with the direction and velocity of hand motion under the assumption of the static prior. As in our previous study ^30^, the plate was lubricated to minimize frictional forces. The ability to adapt to such condition would imply an update of the internal model assuming object stationarity. We assessed the error patterns over trials and determined whether they were consistent with adaptation. In two separate experiments, we either maintained visual feedback about fingertip position relative to the target during the task execution (Experiment 1) or removed it from the PC monitor (Experiment 2). When present, the visual feedback consisted of a 3D-rendered grey dot whose position reflected the fingertip’s true location.

**Figure 1:**
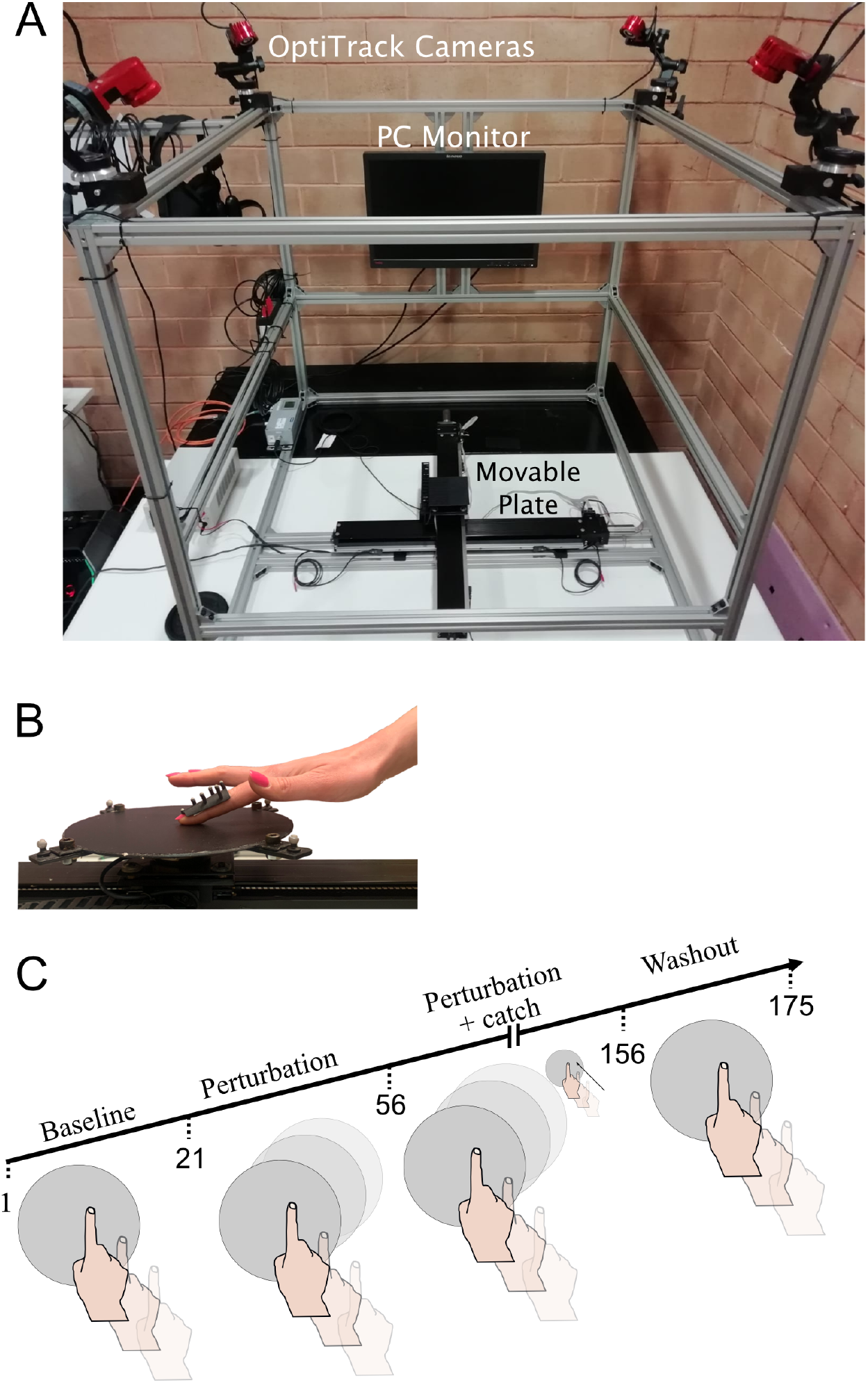
A. The Haptitrack device (^32^; ^33^). A wooden panel with a circular aperture (not shown in figure) prevented participants from seeing the setup during the task. B. Participants wore a thimble with embedded reflective markers while sliding their finger on the contact plate of the Haptitrack. The robotic interface allowed to continuously update the position of the movable plate in real-time based on the participant’s finger position. C. Experimental procedure. The contact plate remained stationary during baseline and washout block and in catch trials interpersed in the last part of the perturbation block. When the perturbation was introduced, the contact plate of the HaptiTrack moved orthogonally to the participant’s finger. Specifically, it followed the finger along the x-axis and moved in the opposite direction along the y-axis.

## 2. Results

Following the method of ^11^, we considered the error at speed peak, defined as the signed perpendicular distance from the finger’s position at maximum velocity to the line connecting the starting and target positions (see Figure 3 and Figure 5). Errors have a negative sign for points below the line and positive for those above. For brevity, we report detailed results only for this measure. Analysis relative to other error measures, namely maximum error, error at the midpoint of the reaching trajectory, and final error of the reaching movement, are reported in the Supplemental information.

### 2.1. Predictions

We expected that positional errors during the baseline trials (where no perturbation was applied) would be randomly distributed around zero, with a sudden shift occurring immediately after the perturbation was introduced. Such systematic error–in accordance with our previous study ^30^–suggests that the change in slip motion at the onset of the perturbation plays a role in controlling the reaching movement, regardless of whether adaptation occurred. Additionally, we hypothesized that distinct error patterns would emerge over the course of the perturbation block (Figure 2 A). Specifically, if adaptation to the tactile slip change occurred, we expected a decrease in error, following the typical exponential decay observed in sensorimotor adaptation paradigms ^34^. Conversely, if no adaptation took place, the error would remain constant throughout the perturbed trials, maintaining the value reached at the onset of the perturbation. While these represent the extreme cases of complete and no adaptation, partial adaptation is also possible, characterized by error reduction followed by a persistent error bias ^18^.

**Figure 2:**
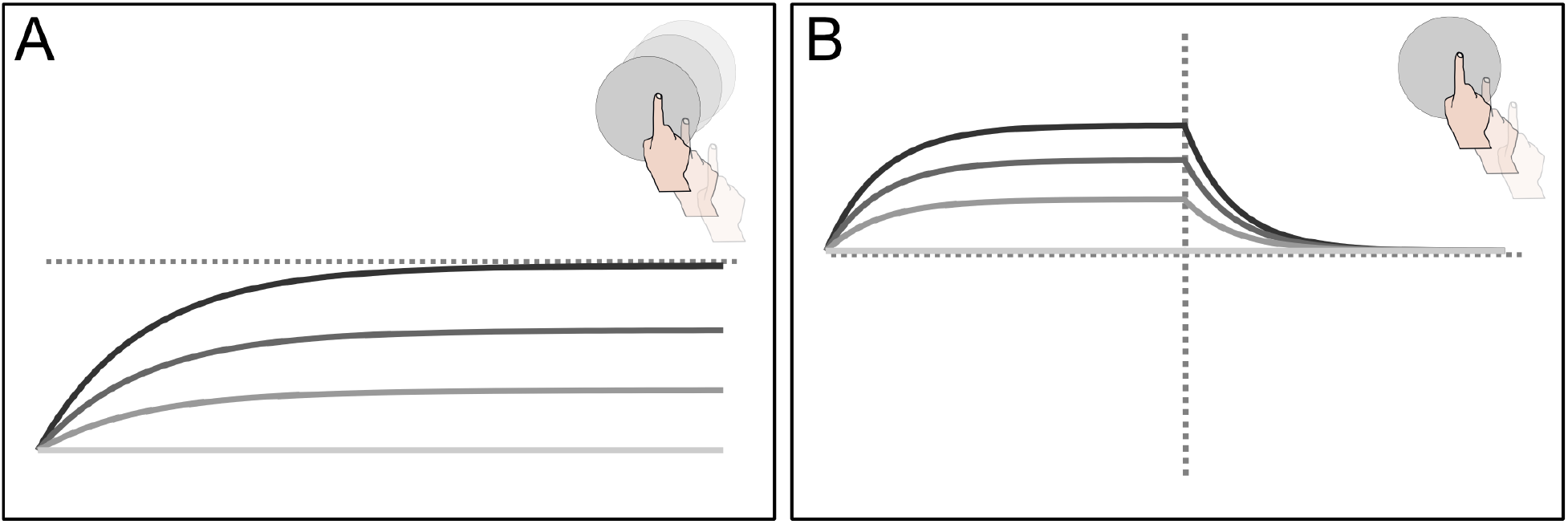
Predicted errors across trials. For errors we intend the deviation of the finger position from the segment connecting starting position and target. Lighter shades of grey represent progressively lower levels of adaptation to the perturbation of the contact plate. A. Errors over trials in the motion perturbation block. B. Errors in catch and washout block.

**Figure 3:**
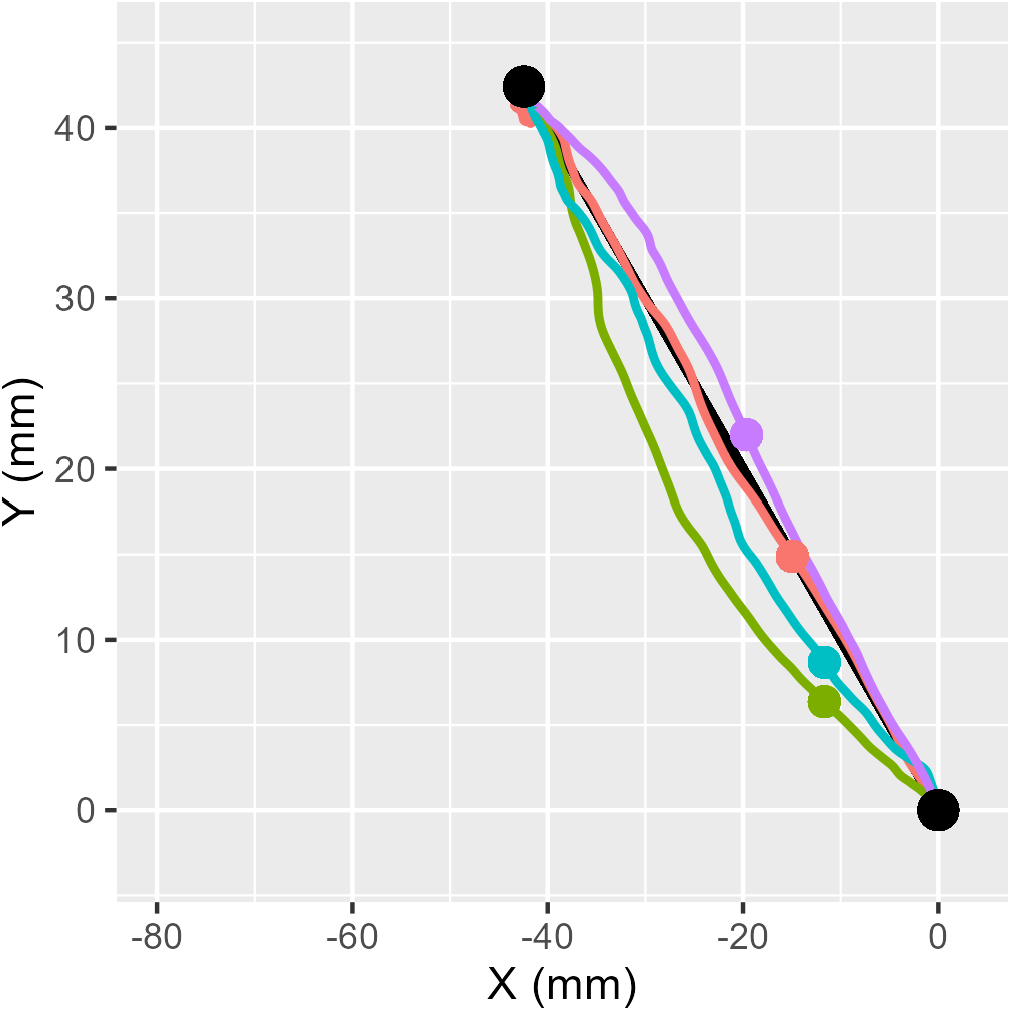
Example of motion paths from different blocks of Experiment 1. The red, green, cyan, and purple trajectories represent trials from the baseline, early adaptation, late adaptation, and early washout blocks, respectively. Colored dots indicate the position were the errors were calculated, corresponding to the peak velocity point. Black dots represent the starting and target positions (lower and upper point, respectively).

In the presence of adaptation, we also expected to observe aftereffects once the perturbation was removed, both during catch trials and in the washout block (Figure 2 B). Specifically, we predicted progressively larger errors in catch trials as adaptation progressed, along with exponential decay of errors returning to baseline levels in the washout block. Conversely, if no adaptation occurred, removing the perturbation would result in errors comparable to those observed during the baseline trials in both catch and washout trials.

To test our predictions, we applied linear mixed models (LMMs) to the stationary sections of the data, as these models are more robust than non-linear models. For further details, readers can consult the Methods section.

### 2.2. Experiment 1

We evaluated whether the motion of the contact plate influenced participants’ reaching errors, in the presence of continuous visual feedback about finger position.

Figure 4 presents the errors averaged across participants over the course of the trials (see also Supplementary Figures 1-3). To assess whether the error patterns aligned with our predictions, we employed LMMs for analysis.

**Figure 4:**
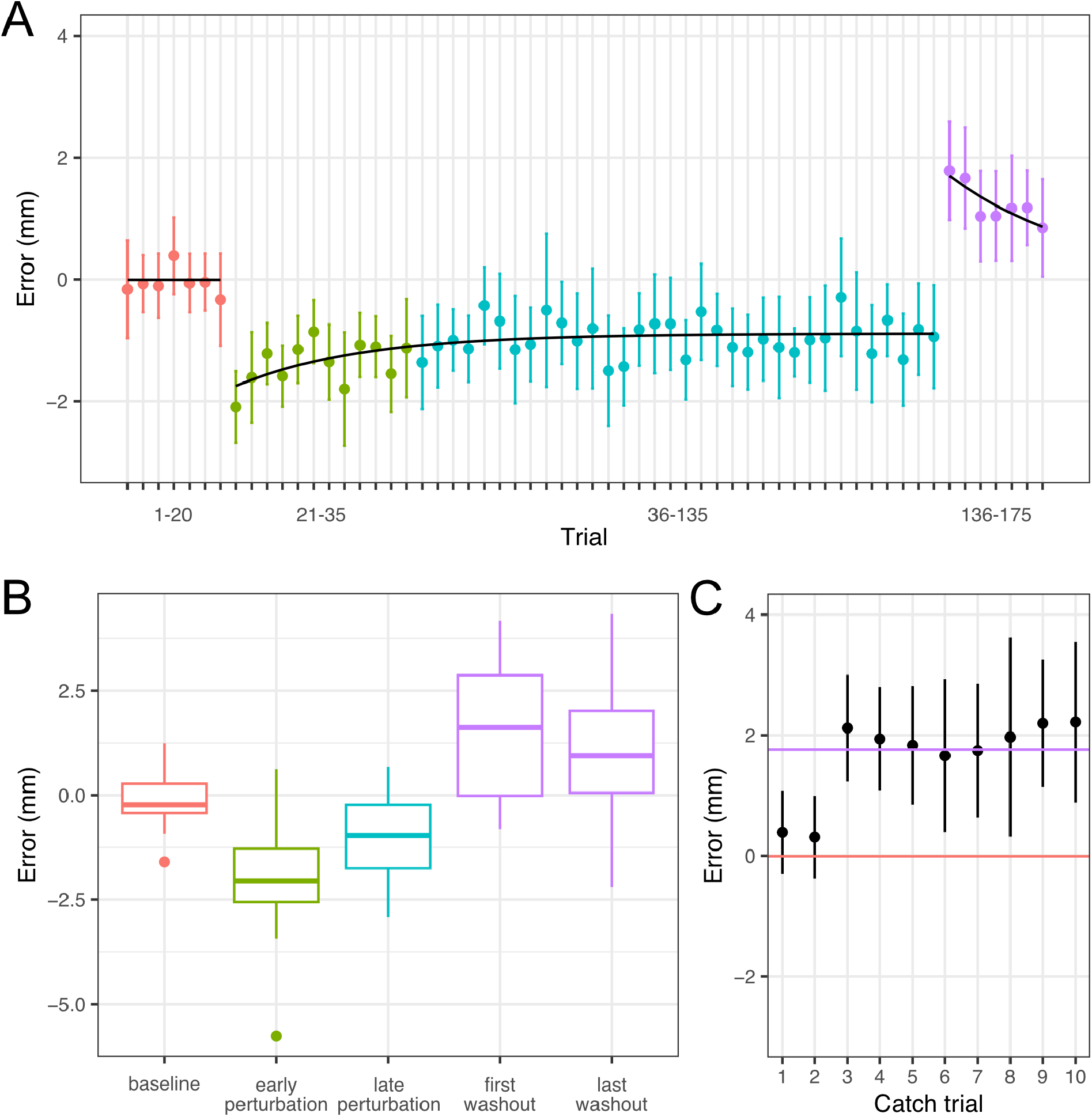
Experiment 1, position error at peak speed across trials (mean and confidence interval). A. Baseline, early perturbation, late perturbation, and washout blocks are represented in red, green, cyan, and purple, respectively. Data are binned by averaging the error across three consecutive trials. The line in the baseline block represents the intercept of the fitted LMM. Binned data in the perturbation block (early and late) were fitted with an exponential function (residual sum of squares (RSS) = 3.847). The asymptote of the function was fixed to the intercept of the LMM fitted to the late perturbation trials. Binned data in the washout block were fitted with an exponential function, with the asymptote fixed to the intercept of the baseline block (RSS = 0.211). B. Error distributions of the blocks used for fitting the models in Equations 2–5. C. Catch trials, ordered progressively from 1 to 10. The red line represents the average error in the baseline block, while the purple line shows the mean error for the first three washout trials.

**Figure 5:**
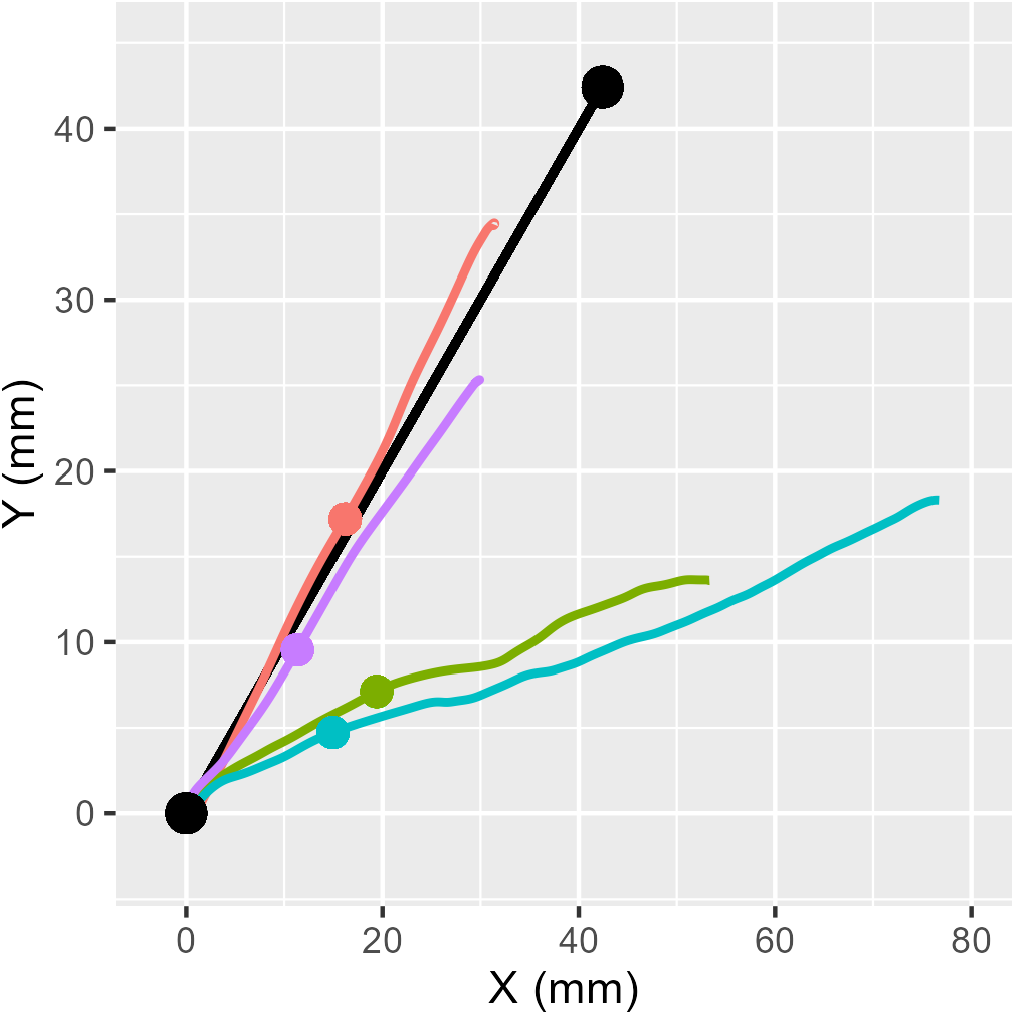
Example of motion paths from different blocks of Experiment 2. Color convention and dot representation are the same as in Figure 3 for Experiment 1.

In the first LMM, we evaluated error as a function of experimental phase and trial number (Equation 2). This model allowed us to test various aspects of our predictions. Specifically, we assessed whether the error was randomly distributed around zero during the baseline trials, which would be indicated by an intercept and slope not statistically different from zero. Additionally, we evaluated whether adaptation was ongoing in the late perturbation trials, which would be reflected in a slope for this experimental phase significantly different from zero. The model also tested whether performance returned to baseline levels by the late perturbation trials, indicating full adaptation. This would manifest as a non-significant change in intercept between baseline and late adaptation trials. For the baseline, the slope and the intercept of the model were non-significanlty different from zero (*β*_0_ ± *SE* (mm) = −0.133 ± 0.256, p = 0.605; *β*_1_ ± *SE* (mm/t) = 0.01 ± 0.019, p = 0.609), consistent with our predictions and suggesting that participants had a solid understanding of the task. A non-significant slope was also observed for the late perturbation trials (*β*_3_ − β_1_ ± SE (mm/t) = −0.008 ± 0.019, p = 0.663), indicating that error reached a plateau during this phase and that no further adaptation occurred. Additionally, the model revealed a significant difference between the intercepts of the baseline and late adaptation trials (*β*_2_ − β_0_ ± SE (mm) = −0.934 ± 0.371, p = 0.014), with a negative intercept in the late adaptation phase. This suggests that participants’ performance remained biased toward the perturbation, and did not return to baseline levels, even after extensive training with the perturbation (Figure 4 A-B, Supplementary Figure 1-2. See Supplementary Tables 1-3 for other error measures).

In a second LMM, we compared the error at the onset of the perturbation–i.e., in the first three trials after the plate perturbation was introduced–with the error in the baseline trials and the last 90 trials of the perturbation blocks (Equation 3). This analysis allowed us to evaluate two key aspects of the error pattern: first, whether the onset of the perturbation was associated with a significant change in error levels, supporting the prediction that the perturbation affected the reaching trajectory. Second, we assessed whether performance remained approximately constant throughout the perturbation trials, indicating the presence or absence of adaptation. Given that the previous model indicated a bias in performance relative to baseline during the late perturbation trials, we expect partial or no adaptation in this phase. The error at the onset of the perturbation increased in absolute value compared to baseline (*δ*_0_ − δ_1_ ± SE (mm) = –1.758 ± 0.295, p = < 0.001). Additionally, error was larger at the beginning of the perturbation block than at the end (*δ*_0_ − δ_2_ ± SE (mm) = –0.871 ± 0.358, p = 0.022), indicating that participants exhibited some level of adaptation (See Supplementary Tables 4-6 for other error measures). Taken together, the two models suggest that adaptation was incomplete, and learning saturated in the presence of tactile slip perturbation.

To evaluate after-effects, we analysed errors in catch trials and in the washout block. Specifically, we evaluated the linear trend in catch trials, ordered progressively from 1 to 10 (Equation 4). The magnitude of after-effect in catch trials serves as an indicator of motor learning ^10,13,11,35^. As such, we expected to observe increasingly large errors during catch trials if learning was ongoing, and relatively constant errors if adaptation was complete (Figure 2). Our results showed that the intercept of the model fitting the sequentially numbered catch trials was significantly greater than zero (*ζ*_0_ ± *SE* (mm) = 0.728 ± 0.34, p = 0.037. Figure 4 C), indicating errors larger than those occurring during baseline and in the opposite direction with respect to the perturbation (see Supplementary Figure 3 and Supplementary Tables 7-9 for other error measures).. The slope was a significant predictor in the model (*ζ*_1_ ± *SE* (mm/t) = 0.157 ± 0.06, p = 0.014). However, this slope was not statistically significant when the first trial was removed from the analysis (p = 0.129), indicating that the first catch trial was associated with a substantially smaller error than all subsequent trials. Together, these results suggest that after-effects were present in the catch trials and that adaptation, although partial, reached a plateau during the late perturbation trials.

To evaluate the after-effects in washout trials, we compared the errors at the beginning and at the end of the block with errors in the baseline block (Equation 5). Error levels differed significantly between baseline and start of washout (*η*_1_ − η_0_ ± SE (mm) = 2.122 ± 0.529, p < 0.001). Similar to the catch trials, the washout errors were in the opposite direction compared to baseline errors, and consistent to error in catch trials (Figure 4 B-C). This result suggests that the internal model used for reaching was recalibrated in response to the perturbation (see Supplementary Tables 10-12 for other error measures). Moreover, errors in the washout block decreased across trials, with the error in the last three trials being closer to baseline levels than in the first three trials (*η*_1_ − η_0_ ± SE (mm) = 0.84 ± 0.405, p = 0.054). In contrast to the perturbation block, the error in the washout phase did not reach a plateau, indicating ongoing re-adaptation (see Supplementary Tables 13-15 for other error measures).

Maximum error and error at the midpoint of the reaching trajectory showed patterns similar to those described for error at peak speed. In contrast, the final position of the reaching trajectory had small, comparable errors across all experimental phases, likely due to the contribution of visual feedback as the fingertip approached the target. These error patterns are consistent with the curved reaching trajectories observed in the experiment (Figure 3 for representative trials).

### 2.3. Experiment 2

In the second experiment, we evaluated errors across trials in the absence of visual feedback. In this condition, errors were generally larger than those observed in Experiment 1, where visual feedback was provided. We focus exclusively on the results related to errors at peak speed, although similar error patterns were noted across other measures, including the signed distance of the endpoint of the trajectory from the target. In Experiment 2, the reaching trajectories did not curve as the finger approached the target position, as illustrated in Figure 3 for Experiment 1. Instead, they followed a straight path that systematically deviated from the segment connecting the starting and target positions (Figure 5). Furthermore, errors were consistently larger than those observed with visual feedback. These overall observations align with previous findings ^30^.

We applied the same models described in Experiment 1 to analyze the data from Experiment 2. For a visual representation of the errors averaged across participants, see Figure 6 (see Supplementary Figures 4-6 for other error measures).

**Figure 6:**
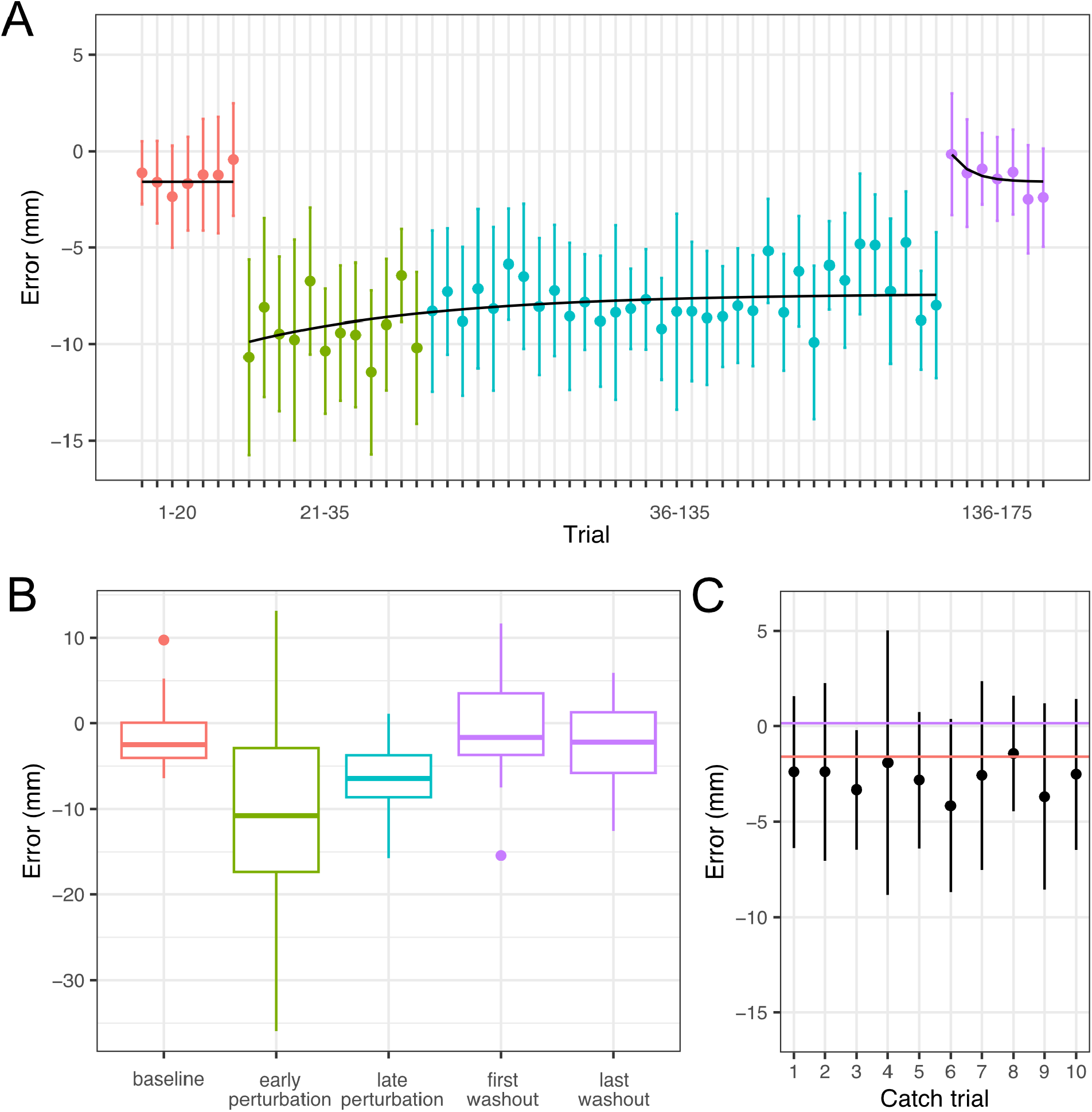
Experiment 2, position error at peak speed across trials (mean and confidence interval). A. Baseline, early perturbation, late perturbation, and washout blocks are represented in red, green, cyan, and purple, respectively. Data are binned by averaging the error across three consecutive trials. The line in the baseline block represents the intercept of the fitted LMM. Binned data in the perturbation block (early and late) were fitted with an exponential function (residual sum of squares (RSS) = 85.354). The asymptote of the function was fixed to the intercept of the LMM fitted to the late perturbation trials. Binned data in the washout block were fitted with an exponential function, with the asymptote fixed to the intercept of the baseline block (RSS = 1.969). B. Error distributions of the blocks used for fitting the models in Equations 2–5. C. Catch trials, ordered progressively from 1 to 10. The red line represents the average error in the baseline block, while the purple line shows the mean error for the first three washout trials.

When considering the model in Equation 2, which compared baseline and late perturbation trials, we found that the baseline errors were randomly distributed around zero, as indicated by the model’s non-significant intercept and slope for the block (*β*_0_ ± *SE* (mm) = −1.547 ± 1.251, p = 0.224; *β*_1_ ± *SE* (mm/t) = −0.001 ± 0.067, p = 0.987). The late perturbation trials had a non-significant slope (*β*_3_ − β_1_ ± SE (mm/t) = 0.016 0.067, p = 0.813) along with a statistically significant difference in intercept compared to the baseline (*β*_2_ − β_0_ ± SE (mm) = −7.593 ± 1.5, p = 5.637 *×* 10^−6^). This indicates that the errors in the late perturbation trials were randomly distributed around a bias, with the bias directed toward the perturbation of the plate (see Supplementary Tables 16-18 for other error measures).

We fitted the model in Equation 3 to assess whether a change in error occurred at the onset of the perturbation, and to determine if there was a decrease in error over the perturbation phase, indicating partial adaptation. While the difference between the baseline and the first perturbation trials was statistically significant (*δ*_0_ − *δ*_1_ ± *SE* (mm) = –8.7 ± 2.016, p = < 0.001), there was no significant difference between early and late perturbation trials (*δ*_0_ − *δ*_2_ ± *SE* (mm) = –2.69 ± 1.743, p = 0.142. See Supplementary Tables 19-21 for other error measures).

When evaluating presence of after-effects in catch trials (Equation 4), we found that both intercept and slope associated with the sequentially numbered trials did not have a significant effect (*ζ*_0_ ± *SE* (mm) = −2.781 ± 1.645, p = 0.111; *ζ*_1_ ± *SE* (mm/t) = 0.031 ± 0.189, p = 0.872, Figure 6 C. See Supplementary Tables 22-24 and Supplementary Figure 6 for other error measures.). This indicates that the errors in all the catch trials were comparable to those observed in the baseline trials. Similarly, there were no significant differences between baseline and washout errors (Equation 5) during both the early and late trials of the washout block (all p > 0.05. See Supplementary Tables 25-30 for other error measures). Collectively, these results do not show evidence of adaptation for Experiment 2.

### 2.4. Explicit assessment of plate motion

At the end of both experiments, we asked participants to reproduce in a drawing the direction in which they felt the plate moving. We only submitted the questionnaires to participants that had no previous knowledge about the functioning principles of the Haptitrack setup. For Experiment 1, 17 participants responded to the questionnaire. 11 participants reported that the plate moved linearly (Figure 7 A), however the direction many of the participants indicated (black arrows in figure) was not consistent with the direction of motion of the plate (red arrows). The remaining participants reported the plate rotating. For Experiment 2, 14 participants responded to the questionnaire. Of them, 12 reported that the contact surface moved linearly (Figure 7 B), 2 reported that it rotated. Moreover, 2 participants declared that the plate rotated during the linear motion. In both experiments, the reported directions for perceived linear motion varied greatly between participants, suggesting that the information on surface motion could not be reliably accessed during self motion.

**Figure 7:**
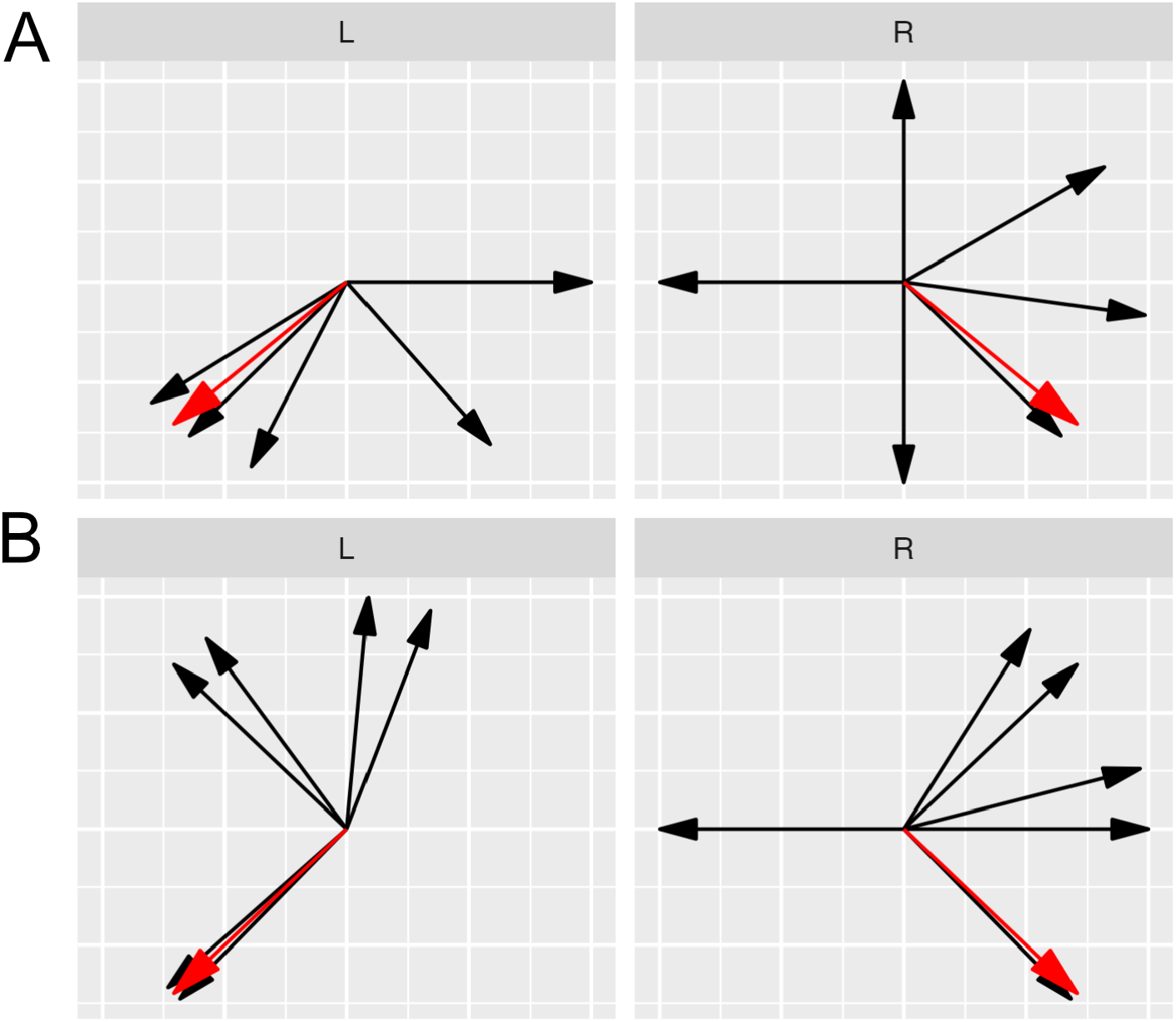
Representation of the directions indicated by participants who reported that the plate moved linearly during perturbed trials (black arrows) in Experiment 1 (A) and 2 (B). In red is the actual motion direction of the plate.

## 3. Discussion

In our study, we investigated whether participants could adapt to perturbations in tactile slip during reaching movements. We observed that prolonged exposure to perturbations in the motion of a contact plate led to limited adaptation in hand reaching movements. Specifically, when visual feedback about finger displacement was provided, we found evidence of partial adaptation. This was characterized by a reduction in error at the onset of the perturbation, a residual bias persisting through the final 70% of the perturbed trials (average compensation, expressed as percentage of ratio between total corrected error and error at perturbation onset: 49.6%, CI[9.5%,76.7%]), and the presence of aftereffects during catch trials and the washout phase of the experimental session. In contrast, when visual feedback was absent, no statistically significant evidence of adaptation was detected. The perturbation of the plate motion consistently induced a systematic error in the reaching trajectory, which remained largely unchanged across the perturbed trials (average compensation 30.9%, CI[-16.2%,57.7%]). Upon removal of the perturbation, the error of the reaching trajectories reverted to those observed during the baseline trials, with no significant evidence of aftereffects.

Incomplete error compensation has been documented in many experimental paradigms exploring sensorimotor adaptation. For example, a residual bias persists despite repeated trials when introducing constant visual shifts ^8^, angular rotations ^36,37^, or dynamical perturbation ^38^. Incomplete adaptation occurs also in saccadic eye movements ^39^ and walking ^40^. The average compensation varies largely among those reported (between ∼33% and ∼67% for adaptation to visuomotor shift in ^8^; between ∼83% and ∼97% for adaptation to visuomotor rotation in ^36^ and ^41^; between ∼60% and ∼80% for force field adaptation in ^9^). One possible explanation for the lack of error compensation in our experiment is that the perceived error is not considered reliable, particularly when visual feedback is missing, as the perturbation violates the strong prior of slow motion. In everyday life situations, such prior implies that tactile and proprioceptive cues are strongly correlated, while in our experiment, they provide conflicting information regarding the motion of the hand relative to a surface. Computational models for adaptation incorporate a forgetting factor ^36,37^ to counterbalance the impact of a learning factor ^42^. The forgetting component of the model is a mechanism enabling conservative responses to transient conditions but effective corrections to more permanent changes^37^. In this perspective, a perturbation violating the static prior such as that under discussion is likely treated very conservatively, so that the residual error is not compensated. Other strong priors, such as the gravitational priors, are associated with a low degree of adaptation when a violation occurs ^18^.

The widely accepted perspective that sensorimotor adaptation involves multiple processes, both implicit and explicit, is well supported in the literature ^42,43,44^. In their study, ^9^ show that residual errors in motor adaptation primarily stem from an implicit learning system adjusting its sensitivity to errors based on the consistency of past occurrences. Notably, no residual errors remain when adaptation rely solely on explicit strategies. In our experiment, it is plausible that participants exclusively employ implicit strategies to correct movement. In fact, regardless of the presence of visual feedback, most participants were unable to accurately identify or describe the motion of the plate, as it emerged from post-experiment questionnaires. Such inability is coherent with previous results ^22^ and with findings in other sensory modalities such as vision, that show a difficulty in the estimation of background motion when self-motion is ongoing.

The manipulation of visual feedback had an important role in our study. While in both experiments adaptation was limited, a stronger adaptation evidence was found when visual feedback was provided. Moreover, the modulation of visual feedback about finger displacement influenced the shape of the reaching trajectories. With visual feedback, errors tended to decrease within a trial as the finger approached the target, resulting in curved trajectories and endpoint errors in perturbed trials compared to those in unperturbed trials. In contrast, without visual feedback, trajectories were straight and exhibited large endpoint errors, as the presence of the plate perturbation led to an error in the trajectory direction that was not compensated within a trial. Curved trajectories are consistently observed in paradigms involving adaptation to visual feedback rotations ^45^ or to external force fields. In the presence of a force field, correction movements occur even in the absence of visual feedback ^10,46^ or proprioceptive information ^47,48^. This suggests that when external forces are applied, the mapping between one’s motion and the relative position of one’s hand with respect to external landmarks (such as the target) is maintained. The mentioned studies typically manipulate visual feedback or limb dynamics to explore the interplay between visual and proprioceptive signals in motor control. These tasks often involve producing ballistic movements or guiding a hand-held manipulandum to a target, excluding frictional forces on the skin and, consequently, neglecting the role of the tactile sensory channel. In contrast, our study introduced perturbations specifically targeting the perception of slip motion at the level of the fingertip to evaluate how mismatches between proprioceptive and tactile signals influence motor control. It must be noticed that tangential forces differ between perturbed and unperturbed trials. However, it is unlikely that adaptation in our task was driven by changes in contact forces associated with the motion of the contact plate. In our previous study ^30^, we found that systematic errors associated with the modulation of tactile slip information were consistent across varying levels of frictional force, but varied significantly under different conditions of tactile sensitivity. Additionally, we observed that force distributions during the reaching task were comparable regardless of the availability of visual feedback. In our current study, we observed curved trajectories only when visual feedback was provided, suggesting that the online recalibration of the reaching movement is mainly driven by visual information about finger displacement.

Previous research emphasize the role of the tactile channel in motor control ^49^. In a recent study, we showed that reaching movements are systematically biased when the motion of the contact surface violates the assumption that inanimate objects are stationary during active motion ^30^. In our current study, the change in tactile slip introduces a systematic bias in the reaching trajectory, in accordance with previous results. We found that such bias cannot be effectively reduced through prolonged exposure to the perturbation. In their study, ^50^ proposed that due to proprioceptive feedback delay, information about limb position might be inaccurate during active motion. In line with this property, we hypothesize that when the perturbation to the contact surface is applied, the information provided by tactile slip is considered more reliable than that provided by proprioceptive signals. As a consequence, participants rely more on the non-veridical slip information.

The assumption that inanimate objects are stationary informs behavior in perceptual and motor tasks, across different sensory channels ^28,51,22,52^. While we demonstrated the robustness of the prior in the control of reaching movements, this may not directly extend to other contexts or tasks. For instance, humans quickly and effectively adapt to motion perturbations when standing or walking over unstable surfaces^53,54,55^. Unexpected shifts of the support platform during stance prompt typical postural adjustments, which are triggered by multisensory stimuli from the visual, vestibular, and somatosensory systems ^56^. Taken together, these findings highlight the critical role of suprasensory cognitive factors in modulating priors in response to context-specific tasks.

A better understanding of the ability of human participants to adapt to a moving device is essential for designing and evaluating innovative technologies like collaborative robots and immersive virtual reality environments. Interfaces leveraging the principles described here offer potential for the assistance and physical rehabilitation of individuals with sensorimotor impairments, such as those affected by stroke or spinal chord injury. Other interesting perspectives include applying our findings in the design of human-like control for soft robotic manipulators, to investigate their capabilities to adapt and react at changing environments^57^, and in applications of extended reality for virtual object manipulation ^31^. By assessing the constraints and limitations involved in these interactions, we can provide elements for evaluating and predicting the ease of learning to use such devices and their effectiveness in real-world applications.

## 4. Methods

### 4.1. Participants

Forty healthy volunteers took part to the study. Twenty participants (7 males and 13 females, age 22.2 ± 1.8 years; 19 right-handed, 1 left-handed according to self-report) took part to experiment 1, twenty (9 males and 11 females, age 20 ± 2 years; 18 right-handed, 2 left-handed according to self-report) to Experiment 2. They all had normal or corrected-to-normal vision. All participants gave their written informed consent to the experiment following guidelines of the Declaration of Helsinki. The study was approved by the local Ethics Committee (comitato etico territoriale Lazio area 5, registro sperimentazioni n. 85/SL/23).

### 4.2. Experimental setup

The experiment was performed using the HaptiTrack, a custom device developed to physically decouple tactile slip motion and hand movements. The device consists of a contact plate moving in 2D thanks to two perpendicular linear motion axes actuated by DC-motors (Figure 1 A). A 6-axis sensor (ATI Mini45) measures contact forces and torques on the plate, while hand and finger tracking is performed by a motion tracking system (OptiTrack system with four Flex13 cameras and OptiHub synchronization box, frame rate 120 Hz; measurement error < 0.1 mm). Each hardware component, and their interaction, are configured using custom-written C++ libraries under 3-clause BSD license. For a more exhaustive description of the HaptiTrack, readers are referred to previous publications ^32,33^.

A 24-inch computer monitor was mounted to the back of the device frame to provide visual feedback, while a wooden panel installed in front occluded the participants’ direct view of both the device and their own hand. Participants touched the contact plate by passing their hand through a circular aperture in the panel, which was covered with black cloth for additional visual occlusion. In order to reduce friction during movement, the contact plate was covered with Teflon and coated with ultrasound gel ^30^. A 3D-printed thimble with embedded reflective markers was used to track finger movements within the workspace of the device (Figure 1 B). The thimble was worn on the right index finger. Finger movement controlled the position of the plate, provided that the force sensor detected a normal force between 0.3 and 2 N. The position of the plate was updated every 5 ms according to the Equation:

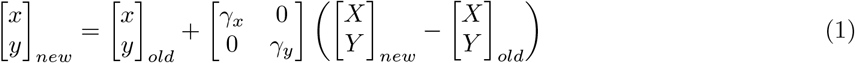

Where lowercase *x* and *y* represent coordinates of the contact plate position; uppercase letters correspond to the finger position coordinates; subscripts *old* and *new* refer to previous and current position, respectively; and *γ*_*x*_, *γ*_*y*_ represent the gains that control the relationship between the participant’s finger position and the contact plate’s movement along each axis. For *γ* = 0, the contact plate remained stationary relative to the finger; For *γ >* 0 the plate moved in the same direction as the finger, and for *γ <* 0 the plate moved in the opposite direction.

### 4.3. Experimental task and procedure

### 4.3.1. Experiment 1: Plate perturbation with visual feedback

Participants were instructed to perform reaching movements on the contact plate of the HaptiTrack towards a virtual target, which consisted of a 3D-rendered green dot, presented on the screen. For each participant, the target was always positioned to 45 degrees either to the left or to the right from straight ahead, at a distance of 60 mm from the starting position. The starting position was represented as a 2D rendered grey dot near the edge of the surface adjacent to the participant’s body midline. Continuous visual feedback about the position of the hand, in the form of a 3D rendered grey dot, was given on the computer monitor in all trials. At the beginning of a session, participants completed a familiarization block consisting of 20 trials, in which the contact plate did not move. Following the familiarization, the experiment consisted of 175 trials divided into 4 blocks: baseline, perturbation, catch, and washout (Figure 1 C).

The baseline block consisted of 20 trials where the plate was stationary (*γ*_*x*_ = *γ*_*y*_ = 0 in Equation 1). In the perturbation block, which included 125 trials, the plate moved with a positive gain on the x-axis (*γ*_*x*_ = 0.7, following movement) and a negative gain on the y-axis (*γ*_*y*_ = −0.7, opposite movement). Given such combination of gains, the plate moved in the direction orthogonal to that of the finger, towards the participant and to the left when they reached the left target, and towards the participant and to the right when they reached the right target. In the late perturbation block (i.e. last 90 trials), 10 catch trials were randomly and uniformly interspersed, in which the contact plate remained stationary. The washout block consisted of 20 trials without plate perturbation. Each experimental session lasted approximately 40 minutes. Participants wore headphones playing pink noise for the entire duration of the task. At the end of the experiment, we asked participants to reproduce in a drawing the perceived motion of the plate. The questionnaire was submitted only to participants who were completely unfamiliar with how the Haptitrack operated.

### 4.3.2. Experiment 2: Contact plate perturbation without visual feedback

The setup and procedure in Experiment 2 were consistent with those described above, with the only difference being the absence of visual feedback during the reaching task. Participants had visual feedback of their finger position while aligning it with the starting point. Once the finger was correctly positioned, the target appeared, and the finger avatar disappeared for the remainder of the trial. In the initial familiarization block, participants completed 10 trials with visual feedback, followed by 10 trials without it. No motion perturbation of the plate occurred during these trials.

### 4.4. Data analysis

We applied filtering to all signals using a Butterworth 5th order filter with a cut-off frequency of 10 Hz. To obtain finger velocity along both the *x* and *y* axis, we calculated the difference between position in two consecutive frames and divided it by the frame duration (5 ms).

We removed individual trials from further analysis if they met the following exclusion criteria: the total time for completing the trajectory was less than 1 s; the projection of the trajectory on the straight line connecting starting and target position did not reach half of the distance between the two points; peak velocity along x and y axis exceeded 3 standard deviations for the mean velocity across all trials. In Experiment 1, we excluded data from a specific participant due to outlier behavior. Specifically, this participant exhibited a significantly higher error in both the baseline and adaptation blocks compared to the error levels observed across all other participants. According to our criteria, we removed 527 trials from data in Experiment 1 (∼15% of total). In experiment 2, we excluded data from one participant due to a hardware malfunction. Additionally, we removed individual trials based on the same inclusion criteria used in Experiment 1, resulting in a total of 606 trials removed (∼18% of total). All signal pre-processing and filtering were implemented using Python version 3.9.2, all subsequent analyses were conducted using R version 4.2.1.

For data analysis we used the Linear Mixed Model framework, which accounts simultaneously for fixed- and random-effects ^58^. Several models were fitted to assess the different error patterns described in the Predictions section.

To analyze errors during the baseline and late perturbation phases, we applied the following model:

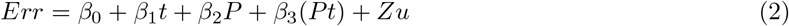

where *β*_0_, …, *β*_3_ are the fixed-effects parameter of the model, *t* is the trial number, *P* is a dummy variable coding for the different experimental phases (*P* = 0 for the baseline, *P* = 1 for the late perturbation, which corresponds to the last 90 trials where the perturbation was applied), and *Zu* incorporates the random-effect predictors in the model.

To evaluate errors during the onset of the perturbation (i.e., the first three trials after the perturbation was applied) compared to the baseline and late perturbation phases, we used the following model:

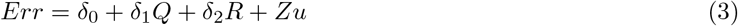

Here, *δ*_0_, …, *δ*_2_ are the fixed-effect parameters of the model, and *Q, R* are the dummy variables coding for the different experimental blocks (*Q* = 1 for baseline and 0 otherwise; *R* = 1 for late perturbation and 0 otherwise).

For catch trials, the model equation was:

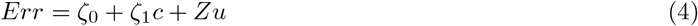

where *ζ*_0_, *ζ*_1_ are the fixed-effects parameters of the model and *c* refers to the catch trials, indexed from 1 to 10.

Finally, we compared errors between the baseline and the first three trials of the washout phase, as well as between the baseline and the last three trials of washout. In both comparisons, the model had the following form:

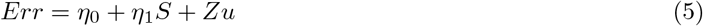

here, *η*_0_ and *η*_1_ represent the fixed-effect parameters of the model, and *S* is the dummy variable coding for the different experiment blocks (*S* = 0 for baseline, *S* = 1 for washout).

#### 4.4.1. Positional error

For statistical analysis, we focused on the positional error (in mm) at peak velocity for each trial. Specifically, positional error was defined as the Euclidean distance between the finger and the segment connecting starting position and target, measured at the moment when the finger reached maximum speed (Figure 3 and Figure 5). Error sign is negative for points below the segment and positive for those above. This is one of the possible metrics used for assessing performance in reaching studies ^11,59,60^. Other frequently used parameters include final positional error ^30^, maximum positional error ^61,9^, or error at a fixed distance ^62^ from movement onset. In our experiments, all these measures produced similar results, except for final positional error. For Experiment 1, final errors were minimal and did not differ significantly across conditions, while in Experiment 2, they aligned with the patterns observed in the other metrics.

## 5. Author Contributions

Conceptualization: P.B, M.B., C.P.R, F.L., A.M., Data Curation: P.B., C.P.R., G.D., A.F., F.V., Formal Analysis: P.B, C.P.R, G.D., A.F., F.V., A.M., Funding Acquisition: M.B., F.L., A.M., Investigation: P.B., C.P.R., G.D., A.F., F.V., Methodology: P.B, M.B., C.P.R, A.M., Project Administration: A.M, Resources: A.M., Software: P.B., Supervision: P.B., A.M., Validation: M.B., F.L., A.M., Visualization: P.B., C.P.R., A.M., Writing - Original Draft Preparation: P.B., Writing - Review and Editing: P.B, M.B., C.P.R, G.D., A.F., F.V., F.L., A.M.

## 6. Acknowledgments

This study was partially supported by the Italian Ministry of Health (Ricerca corrente, IRCCS Fondazione Santa Lucia), the European Commission H2020 Framework Programme (HARIA, grant no. 101070292), the Italian Ministry of University and Research (MUR) (PRIN grant 202249C5XL and Fondo Italiano per la Scienza (FIS), grant PERCEIVING no. FIS00001153), Next Generation EU Project (NGEU) National Recovery and Resilience Plan (NRRP), project MNESYS (PE0000006) and – A Multiscale integrated approach to the study of the nervous system in health and disease (DN. 1553 11.10.2022) Spoke 1.

The authors declare no competing interests.

