## Supplemental materials for "Incomplete adaptation to surface movement during hand reaching"

We present results for models in Eq. 2-5 described in the main text, applied to the following error measures: maximum error, error at the midpoint of the trajectory (referred to as midpoint error), and error at the end of the trajectory (referred to as final error). In all cases, errors have a negative sign for points below the line connecting starting and target position, and a positive sign for those above.

### Experiment 1

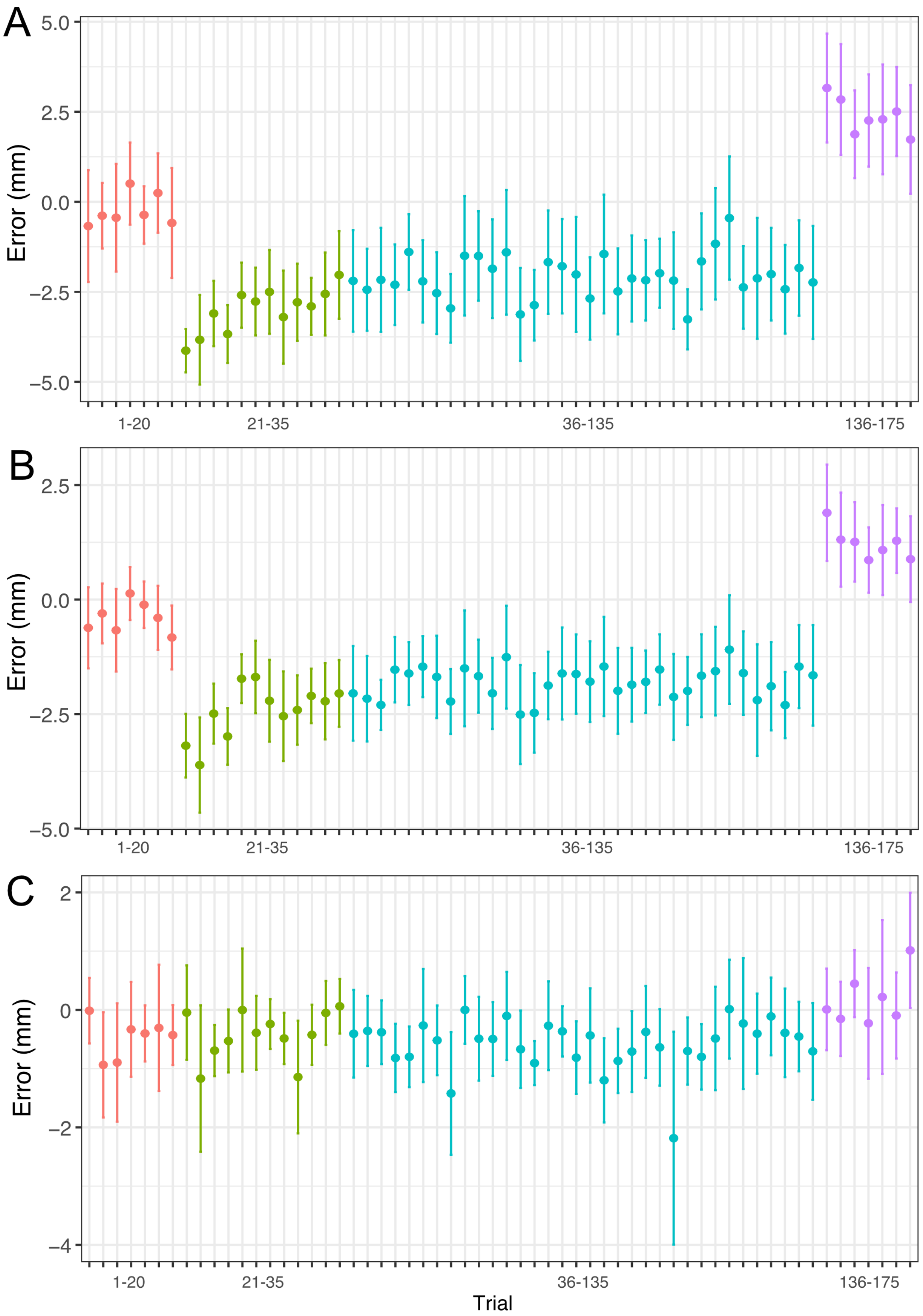


Supplementary Figure 1: Experiment 1, error over trials. Baseline, early perturbation, late perturbation, and washout blocks are represented in red, green, cyan, and purple, respectively. Data are binned by averaging the error across three consecutive trials. A. Maximum error. B. Midpoint error. C. Final error.

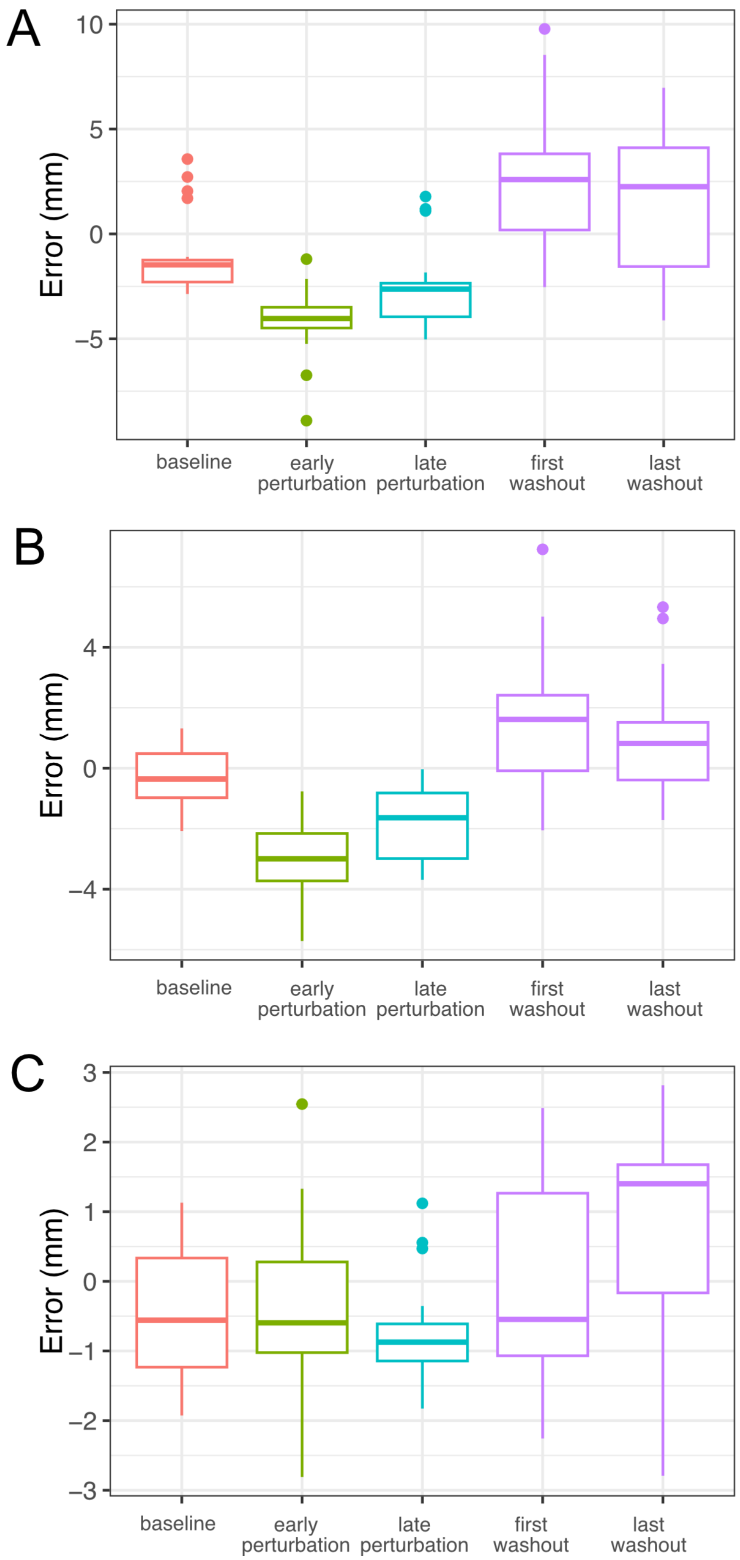


Supplementary Figure 2: Experiment 1, error distributions of the blocks used for fitting the models in Equations 2–5. Baseline, early perturbation, late perturbation, and washout blocks are represented in red, green, cyan, and purple, respectively. A. Maximum error. B. Midpoint error. C. Final error.

### Model Eq. 2: Compare baseline and late perturbation trials

| | Estimate | Std. Error | df | t value | Pr(>\|t\|) |
|---|---|---|---|---|---|
| $\beta_0$ | -0.592 | 0.518 | 72.327 | -1.144 | 0.257 |
| $\beta_1$ | 0.041 | 0.035 | 1849.118 | 1.200 | 0.230 |
| $\beta_2 - \beta_0$ | -1.933 | 0.657 | 86.095 | -2.944 | 0.004 |
| $\beta_3 - \beta_1$ | -0.037 | 0.035 | 1849.065 | -1.058 | 0.290 |

Supplementary Table 1: Summary of model in Eq. 2 for maximum error, experiment 1.

| | Estimate | Std. Error | df | t value | Pr(>\|t\|) |
|---|---|---|---|---|---|
| $\beta_0$ | -0.392 | 0.329 | 63.896 | -1.192 | 0.238 |
| $\beta_1$ | 0.001 | 0.021 | 1847.673 | 0.066 | 0.947 |
| $\beta_2 - \beta_0$ | -1.627 | 0.445 | 58.974 | -3.656 | 0.001 |
| $\beta_3 - \beta_1$ | 0.001 | 0.021 | 1847.624 | 0.043 | 0.966 |

Supplementary Table 2: Summary of model in Eq. 2 for midpoint error, experiment 1.

| | Estimate | Std. Error | df | t value | Pr(>\|t\|) |
|---|---|---|---|---|---|
| $\beta_0$ | -0.537 | 0.348 | 82.515 | -1.542 | 0.127 |
| $\beta_1$ | 0.005 | 0.024 | 1852.325 | 0.187 | 0.851 |
| $\beta_2 - \beta_0$ | 0.009 | 0.396 | 234.078 | 0.022 | 0.983 |
| $\beta_3 - \beta_1$ | -0.005 | 0.024 | 1852.282 | -0.205 | 0.838 |

Supplementary Table 3: Summary of model in Eq. 2 for final error, experiment 1.

### Model Eq. 3: Compare first perturbation trials with baseline, late perturbation trials

| | Estimate | Std. Error | df | t value | Pr(>\|t\|) |
|---|---|---|---|---|---|
| $\delta_0$ | -4.116 | 0.505 | 178.011 | -8.149 | 0.000 |
| $\delta_1 - \delta_0$ | 3.963 | 0.631 | 29.656 | 6.278 | 0.000 |
| $\delta_2 - \delta_0$ | 2.093 | 0.712 | 23.165 | 2.939 | 0.007 |

Supplementary Table 4: Summary of model in Eq. 3 for maximum error, experiment 1.

| | Estimate | Std. Error | df | t value | Pr(>\|t\|) |
|---|---|---|---|---|---|
| $\delta_0$ | -3.268 | 0.317 | 19.430 | -10.302 | 0.000 |
| $\delta_1 - \delta_0$ | 2.892 | 0.428 | 18.012 | 6.764 | 0.000 |
| $\delta_2 - \delta_0$ | 1.493 | 0.428 | 18.585 | 3.487 | 0.003 |

Supplementary Table 5: Summary of model in Eq. 3 for midpoint error, experiment 1.

| | Estimate | Std. Error | df | t value | Pr(>\|t\|) |
|---|---|---|---|---|---|
| $\delta_0$ | -0.575 | 0.482 | 18.014 | -1.193 | 0.248 |
| $\delta_1 - \delta_0$ | 0.091 | 0.405 | 39.834 | 0.225 | 0.823 |
| $\delta_2 - \delta_0$ | 0.000 | 0.443 | 18.445 | 0.001 | 0.999 |

Supplementary Table 6: Summary of model in Eq. 3 for final error, experiment 1.

### Model Eq. 4: Catch trials

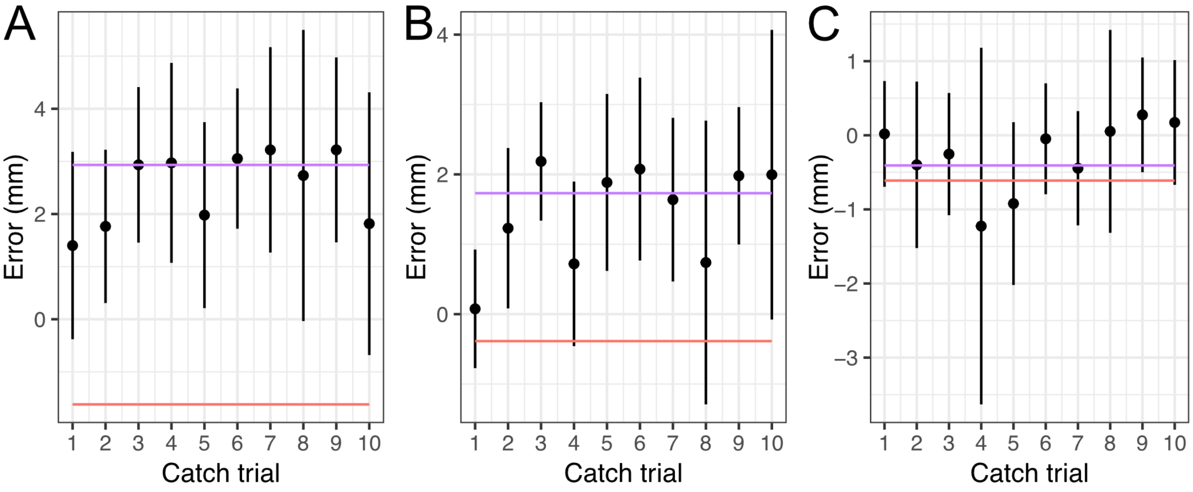


Supplementary Figure 3: Catch trials, ordered progressively from 1 to 10. The red line represents the average error in the baseline block, while the purple line shows the mean error for the first three washout trials. A. Maximum error. B. Midpoint error. C. Final error.

| | Estimate | Std. Error | df | t value | Pr(>\|t\|) |
|---|---|---|---|---|---|
| $\zeta_0$ | 1.998 | 0.736 | 18.294 | 2.715 | 0.014 |
| $\zeta_1$ | 0.082 | 0.126 | 17.487 | 0.651 | 0.524 |

Supplementary Table 7: Summary of model in Eq. 4 for maximum error, experiment 1.

| | Estimate | Std. Error | df | t value | Pr(>\|t\|) |
|---|---|---|---|---|---|
| $\zeta_0$ | 0.862 | 0.427 | 18.377 | 2.018 | 0.058 |
| $\zeta_1$ | 0.095 | 0.076 | 17.653 | 1.255 | 0.226 |

Supplementary Table 8: Summary of model in Eq. 4 for midpoint error, experiment 1.

| | Estimate | Std. Error | df | t value | Pr(>\|t\|) |
|---|---|---|---|---|---|
| $\zeta_0$ | -0.650 | 0.455 | 28.014 | -1.429 | 0.164 |
| $\zeta_1$ | 0.065 | 0.066 | 99.261 | 0.976 | 0.332 |

Supplementary Table 9: Summary of model in Eq. 4 for final error, experiment 1.

### Model Eq. 5: Compare baseline and early washout trials

| | Estimate | Std. Error | df | t value | Pr(>\|t\|) |
|---|---|---|---|---|---|
| $\eta_0$ | -0.154 | 0.368 | 18.258 | -0.420 | 0.679 |
| $\eta_1 - \eta_0$ | 3.740 | 0.944 | 18.042 | 3.962 | 0.001 |

Supplementary Table 10: Summary of model in Eq. 5 for maximum error (early washout trials), experiment 1.

| | Estimate | Std. Error | df | t value | Pr(>\|t\|) |
|---|---|---|---|---|---|
| $\eta_0$ | -0.374 | 0.239 | 18.168 | -1.564 | 0.135 |
| $\eta_1 - \eta_0$ | 2.583 | 0.546 | 17.981 | 4.728 | 0.000 |

Supplementary Table 11: Summary of model in Eq. 5 for midpoint error (early washout trials), experiment 1.

| | Estimate | Std. Error | df | t value | Pr(>\|t\|) |
|---|---|---|---|---|---|
| $\eta_0$ | -0.474 | 0.234 | 17.559 | -2.026 | 0.058 |
| $\eta_1 - \eta_0$ | 0.566 | 0.665 | 18.070 | 0.851 | 0.406 |

Supplementary Table 12: Summary of model in Eq. 5 for final error (early washout trials), experiment 1.

#### Model Eq. 5: Compare baseline and late washout trials

| | Estimate | Std. Error | df | t value | Pr(>\|t\|) |
|---|---|---|---|---|---|
| $\eta_0$ | -0.154 | 0.367 | 18.255 | -0.419 | 0.680 |
| $\eta_1 - \eta_0$ | 1.826 | 0.732 | 16.211 | 2.495 | 0.024 |

Supplementary Table 13: Summary of model in Eq. 5 for maximum error (late washout trials), experiment 1.

| | Estimate | Std. Error | df | t value | Pr(>\|t\|) |
|---|---|---|---|---|---|
| $\eta_0$ | -0.375 | 0.239 | 18.160 | -1.573 | 0.133 |
| $\eta_1 - \eta_0$ | 1.235 | 0.412 | 17.373 | 3.002 | 0.008 |

Supplementary Table 14: Summary of model in Eq. 5 for midpoint error (late washout trials), experiment 1.

| | Estimate | Std. Error | df | t value | Pr(>\|t\|) |
|---|---|---|---|---|---|
| $\eta_0$ | -0.474 | 0.234 | 17.543 | -2.025 | 0.058 |
| $\eta_1 - \eta_0$ | 1.494 | 0.602 | 19.054 | 2.481 | 0.023 |

Supplementary Table 15: Summary of model in Eq. 5 for final error (late washout trials), experiment 1.

### Experiment 2

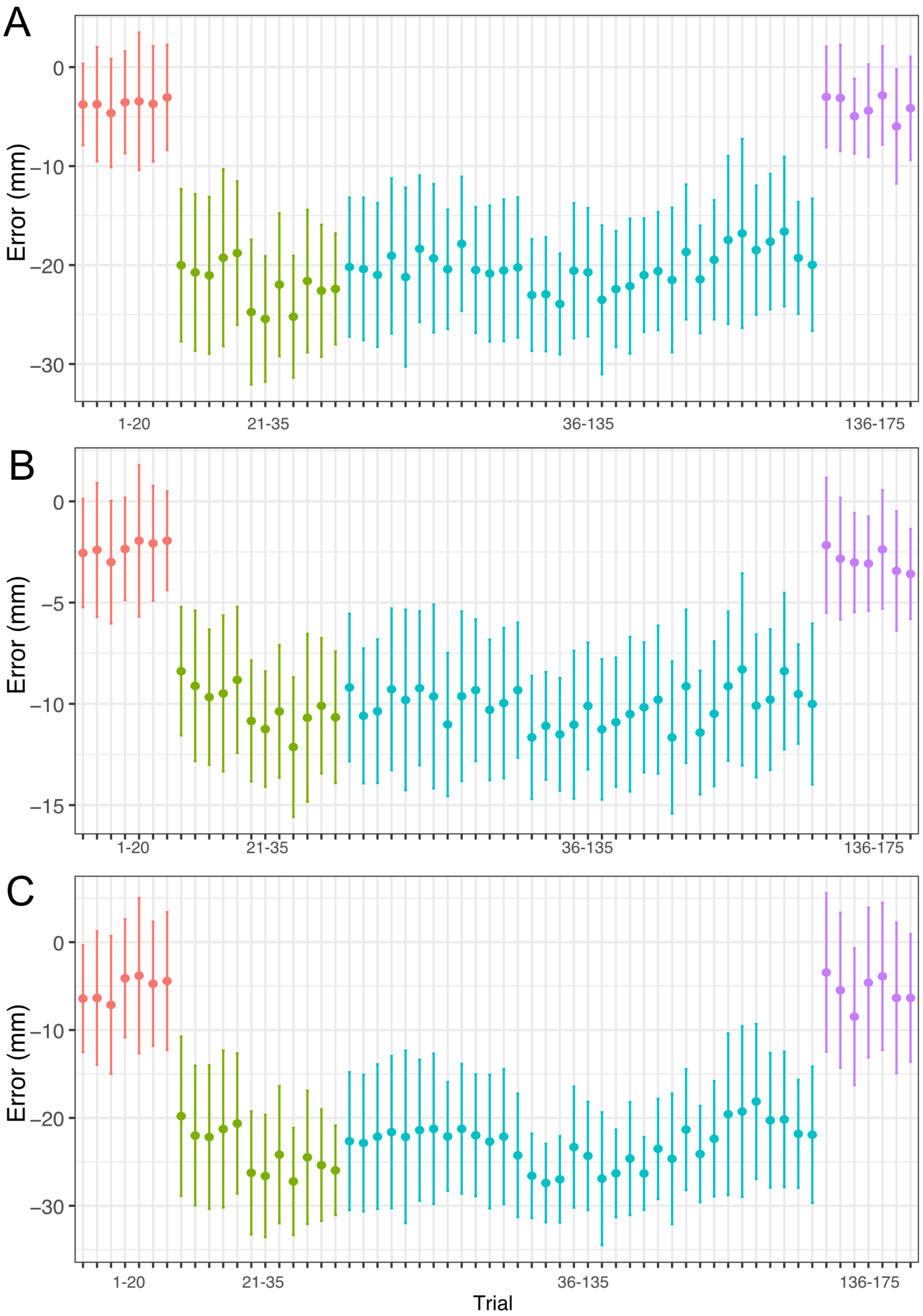


Supplementary Figure 4: Experiment 2, error over trials. Baseline, early perturbation, late perturbation, and washout blocks are represented in red, green, cyan, and purple, respectively. Data are binned by averaging the error across three consecutive trials. A. Maximum error. B. Midpoint error. C. Final error.

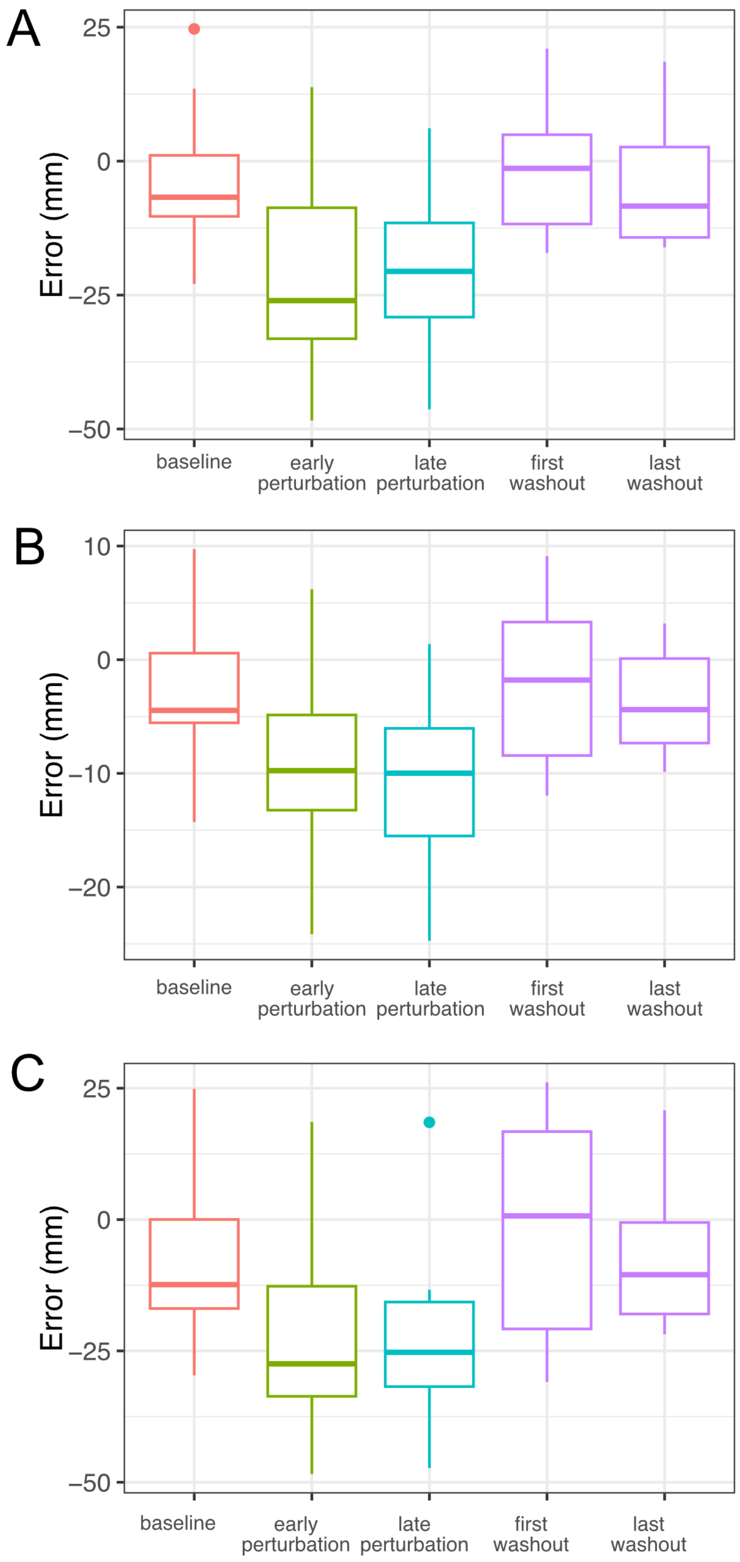


Supplementary Figure 5: Experiment 2, error distributions of the blocks used for fitting the models in Equations 2–5. Baseline, early perturbation, late perturbation, and washout blocks are represented in red, green, cyan, and purple, respectively. A. Maximum error. B. Midpoint error. C. Final error.

### Model Eq. 2: Compare baseline and late perturbation trials

| | Estimate | Std. Error | df | t value | Pr(>\|t\|) |
|---|---|---|---|---|---|
| $\beta_0$ | -3.545 | 2.635 | 22.489 | -1.345 | 0.192 |
| $\beta_1$ | -0.029 | 0.084 | 1703.312 | -0.346 | 0.730 |
| $\beta_2 - \beta_0$ | -19.043 | 3.105 | 24.934 | -6.134 | 0.000 |
| $\beta_3 - \beta_1$ | 0.052 | 0.084 | 1703.288 | 0.616 | 0.538 |

Supplementary Table 16: Summary of model in Eq. 2 for maximum error, experiment 2.

| | Estimate | Std. Error | df | t value | Pr(>\|t\|) |
|---|---|---|---|---|---|
| $\beta_0$ | -2.493 | 1.414 | 21.806 | -1.763 | 0.092 |
| $\beta_1$ | 0.008 | 0.042 | 1703.070 | 0.187 | 0.852 |
| $\beta_2 - \beta_0$ | -8.064 | 1.691 | 23.545 | -4.770 | 0.000 |
| $\beta_3 - \beta_1$ | -0.003 | 0.042 | 1703.050 | -0.078 | 0.938 |

Supplementary Table 17: Summary of model in Eq. 2 for midpoint error, experiment 2.

| | Estimate | Std. Error | df | t value | Pr(>\|t\|) |
|---|---|---|---|---|---|
| $\beta_0$ | -6.485 | 3.517 | 21.382 | -1.844 | 0.079 |
| $\beta_1$ | 0.099 | 0.098 | 1703.012 | 1.010 | 0.313 |
| $\beta_2 - \beta_0$ | -18.279 | 3.712 | 24.713 | -4.925 | 0.000 |
| $\beta_3 - \beta_1$ | -0.083 | 0.099 | 1702.992 | -0.834 | 0.404 |

Supplementary Table 18: Summary of model in Eq. 2 for final error, experiment 2.

### Model Eq. 3: Compare first perturbation trials with baseline, late perturbation trials

| | Estimate | Std. Error | df | t value | Pr(>\|t\|) |
|---|---|---|---|---|---|
| $\delta_0$ | -20.272 | 3.771 | 18.007 | -5.376 | 0.000 |
| $\delta_1 - \delta_0$ | 16.432 | 2.900 | 16.996 | 5.667 | 0.000 |
| $\delta_2 - \delta_0$ | 0.098 | 2.506 | 19.250 | 0.039 | 0.969 |

Supplementary Table 19: Summary of model in Eq. 3 for maximum error, experiment 2.

| | Estimate | Std. Error | df | t value | Pr(>\|t\|) |
|---|---|---|---|---|---|
| $\delta_0$ | -8.652 | 1.606 | 18.123 | -5.389 | 0.000 |
| $\delta_1 - \delta_0$ | 6.242 | 1.115 | 15.501 | 5.598 | 0.000 |
| $\delta_2 - \delta_0$ | -1.423 | 1.237 | 19.295 | -1.151 | 0.264 |

Supplementary Table 20: Summary of model in Eq. 3 for midpoint error, experiment 2.

| | Estimate | Std. Error | df | t value | Pr(>\|t\|) |
|---|---|---|---|---|---|
| $\delta_0$ | -20.530 | 4.058 | 18.073 | -5.059 | 0.000 |
| $\delta_1 - \delta_0$ | 15.073 | 3.168 | 16.700 | 4.757 | 0.000 |
| $\delta_2 - \delta_0$ | -2.447 | 3.098 | 19.140 | -0.790 | 0.439 |

Supplementary Table 21: Summary of model in Eq. 3 for final error, experiment 2.

**Model Eq. 4: Catch trials**

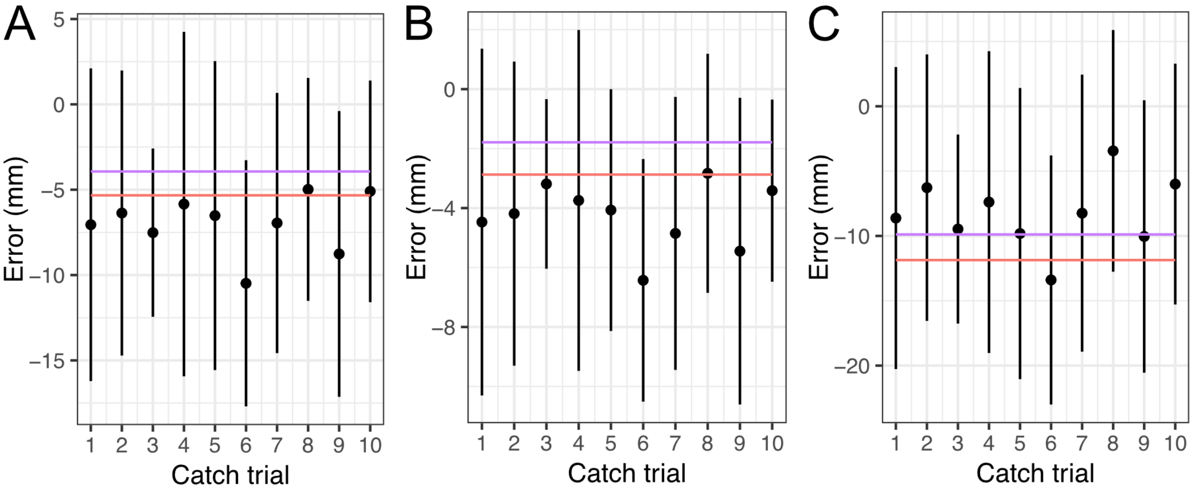


Supplementary Figure 6: Catch trials, ordered progressively from 1 to 10. The red line represents the average error in the baseline block, while the purple line shows the mean error for the first three washout trials. A. Maximum error. B. Midpoint error. C. Final error.

| | Estimate | Std. Error | df | t value | Pr(>\|t\|) |
|---|---|---|---|---|---|
| $\zeta_0$ | -7.218 | 4.070 | 15.358 | -1.774 | 0.096 |
| $\zeta_1$ | 0.112 | 0.388 | 17.448 | 0.289 | 0.776 |

Supplementary Table 22: Summary of model in Eq. 4 for maximum error, experiment 2.

| | Estimate | Std. Error | df | t value | Pr(>\|t\|) |
|---|---|---|---|---|---|
| $\zeta_0$ | -4.424 | 2.312 | 16.637 | -1.913 | 0.073 |
| $\zeta_1$ | 0.035 | 0.199 | 18.928 | 0.176 | 0.862 |

Supplementary Table 23: Summary of model in Eq. 4 for midpoint error, experiment 2.

| | Estimate | Std. Error | df | t value | Pr(>\|t\|) |
|---|---|---|---|---|---|
| $\zeta_0$ | -9.451 | 4.804 | 16.206 | -1.967 | 0.067 |
| $\zeta_1$ | 0.229 | 0.506 | 18.818 | 0.452 | 0.656 |

Supplementary Table 24: Summary of model in Eq. 4 for final error, experiment 2.

### Model Eq. 5: Compare baseline and early washout trials

| | Estimate | Std. Error | df | t value | Pr(>\|t\|) |
|---|---|---|---|---|---|
| $\eta_0$ | -3.847 | 2.492 | 17.972 | -1.544 | 0.140 |
| $\eta_1 - \eta_0$ | -3.354 | 2.916 | 12.476 | -1.150 | 0.272 |

Supplementary Table 25: Summary of model in Eq. 5 for maximum error (early washout trials), experiment 2.

| | Estimate | Std. Error | df | t value | Pr(>\|t\|) |
|---|---|---|---|---|---|
| $\eta_0$ | -2.415 | 1.347 | 17.979 | -1.793 | 0.090 |
| $\eta_1 - \eta_0$ | -2.036 | 2.061 | 12.032 | -0.988 | 0.343 |

Supplementary Table 26: Summary of model in Eq. 5 for midpoint error (first washout trials), experiment 2.

| | Estimate | Std. Error | df | t value | Pr(>\|t\|) |
|---|---|---|---|---|---|
| $\eta_0$ | -5.470 | 3.364 | 17.992 | -1.626 | 0.121 |
| $\eta_1 - \eta_0$ | -4.656 | 4.609 | 12.906 | -1.010 | 0.331 |

Supplementary Table 27: Summary of model in Eq. 5 for final error (first washout trials), experiment 2.

### Model Eq. 5: Compare baseline and late washout trials

| | Estimate | Std. Error | df | t value | Pr(>\|t\|) |
|---|---|---|---|---|---|
| $\eta_0$ | -3.842 | 2.491 | 17.972 | -1.542 | 0.141 |
| $\eta_1 - \eta_0$ | -0.568 | 3.137 | 17.035 | -0.181 | 0.859 |

Supplementary Table 28: Summary of model in Eq. 5 for maximum error (last washout trials), experiment 2.

| | Estimate | Std. Error | df | t value | Pr(>\|t\|) |
|---|---|---|---|---|---|
| $\eta_0$ | -2.413 | 1.347 | 17.979 | -1.791 | 0.090 |
| $\eta_1 - \eta_0$ | -1.377 | 1.351 | 17.238 | -1.019 | 0.322 |

Supplementary Table 29: Summary of model in Eq. 5 for midpoint error (last washout trials), experiment 2.

| | Estimate | Std. Error | df | t value | Pr(>\|t\|) |
|---|---|---|---|---|---|
| $\eta_0$ | -5.463 | 3.363 | 17.993 | -1.624 | 0.122 |
| $\eta_1 - \eta_0$ | -1.416 | 4.324 | 16.815 | -0.327 | 0.747 |

Supplementary Table 30: Summary of model in Eq. 5 for final error (last washout trials), experiment 2.
